# Characterization of gut microbiome and metabolome in *Helicobacter pylori* patients in an underprivileged community in the United States

**DOI:** 10.1101/2021.05.23.445270

**Authors:** Brian E. White, John D. Sterrett, Zoya Grigoryan, Lauren T. Lally, Jared D. Heinze, Hyder Alikhan, Christopher A. Lowry, Lark Perez, Joshua DeSipio, Sangita Phadtare

## Abstract

**Background:** *Helicobacter pylori*, a bacterium that infects approximately half of the world’s population, is associated with various gastrointestinal diseases, including peptic ulcers, non-ulcer dyspepsia, gastric adenocarcinoma, and gastric lymphoma. To combat the increasing antibiotic resistance of *H. pylori*, the need for new therapeutic strategies has become more pressing. Characterization of the interactions between *H. pylori* and the fecal microbiome, as well as the mechanisms underlying these interactions, may offer new therapeutic approaches. Exploration of changes in fatty acid metabolism associated with *H. pylori*-mediated alterations of the fecal microbiome may also reveal strategies to help prevent progression to neoplasia.

**Aim:** To characterize the gut microbiome and metabolome in *H. pylori* patients in a socioeconomically challenged and underprivileged inner-city community.

**Methods:** Stool samples from 19 *H. pylori* patients and 16 control subjects were analyzed. 16S rRNA gene sequencing was performed on normalized pooled amplicons using the Illumina MiSeq System using a MiSeq reagent kit v2. Alpha and beta diversity analyses were performed in QIIME 2. Non-targeted fatty acid analysis of the samples was carried out using gas chromatography-mass spectrometry (GC-MS), which measures the total content of 30 fatty acids in stool after conversion into their corresponding fatty acid methyl esters. Multi-dimensional scaling (MDS) was performed on Bray-Curtis distance matrices created from both the metabolomics and microbiome datasets and a Procrustes test was performed on the metabolomics and microbiome MDS coordinates.

**Results:** Fecal microbiome analysis showed that alpha diversity was lowest in *H. pylori* patients over 40 years of age compared to control subjects of similar age group. Beta diversity analysis of the samples revealed significant differences in microbial community structure between *H. pylori* patients and control subjects. Thirty-eight and six taxa had lower and higher relative abundance in *H. pylori* patients, respectively. Taxa that were enriched in *H. pylori* patients included *Atopobium*, Gemellaceae, Micrococcaceae, Gemellales and *Rothia* (*R. mucilaginosa*). Notably, relative abundance of the phylum Verrucomicrobia was decreased in *H. pylori* patients compared to control subjects, suggesting disruption of the gut mucosal environment by *H. pylori*. Procrustes analysis showed a significant relationship between the microbiome and metabolome datasets. Stool samples from *H. pylori* patients showed increases in several fatty acids including the polyunsaturated fatty acids (PUFAs) 22:4n6, 22:5n3, 20:3n6 and 22:2n6, while decreases were noted in other fatty acids including the PUFA 18:3n6. The pattern of changes in fatty acid concentration correlated to the Bacteroidetes:Firmicutes ratio determined by 16S rRNA gene analysis.

**Conclusion:** An individualized understanding of gut microbiome features among *H. pylori* patients will pave the way for improved community impact, reduced healthcare burdens of repeated treatment, and decreased mounting resistance.

## INTRODUCTION

*Helicobacter pylori*, a bacterium found in the stomach of roughly half of the world’s population, is associated with various gastrointestinal diseases, including peptic ulcers, non-ulcer dyspepsia, gastric adenocarcinoma, and gastric lymphoma. Although many infections are asymptomatic, eradication of *H. pylori* has been shown to reduce the incidence of gastric cancer and is thus universally recommended^[1]^. For decades, clarithromycin-based triple therapy was considered the first-line therapy for *H. pylori* eradication and associated gastric cancer prevention. Following the global trend in emerging antibiotic resistance, rates of clarithromycin, metronidazole, and levofloxacin resistance among *H. pylori* strains have increased markedly, causing an alarmingly high rate of eradication failure^[2]^. In the absence of universally available antibiotic susceptibility testing, current guidelines advocate for the use of empiric quadruple therapy strategies; however, the efficacy of these regimens may be limited by dual resistance, local availability, and patient adherence^[3]^.

As *H. pylori* antibiotic resistance rises, the need for new therapeutic strategies becomes more pressing. Alternative antibiotics, novel acid suppressants, and vaccination have shown some promise in addressing this challenge^[4–7]^. Probiotics have also garnered attention for a possible role in *H. pylori* treatment, as infection with the bacterium may be associated with gastric dysbiosis, which can promote inflammation, metabolic disease, and carcinogenesis^[8–12]^. With the advent of high throughput 16S rRNA gene sequencing methods and open-source software for subsequent analysis and visualization, researchers have been able to characterize changes in the microbial diversity and community structure of the gut microbiota, as well as specific taxa that are differentially abundant in association with particular phenotypes^[13]^. Many studies have focused on *H. pylori*-mediated changes to the gastric microbiome, although data have demonstrated variable patterns of effects on multiple metrics of microbial diversity and community structure within the gastric microbiome^[14]^. Given the challenges associated with obtaining gastric samples, researchers have also focused on the impact of *H. pylori* infection on distal gut, *i.e.*, fecal, microbiota, and observed changes in the fecal microbiome have prompted speculation that such alterations may be involved in the development of gastrointestinal lesions and cancer^[15]^. Further characterization of the interactions between *H. pylori* and the fecal microbiome, as well as the mechanisms underlying these interactions, may lead to novel therapeutic opportunities, as well as to identification of patients at risk for progression to particular clinical manifestations.

Given the association between *H. pylori* infection and gastrointestinal inflammation, attention has also been paid to compounds with anti-inflammatory activity, including fatty acids. Polyunsaturated fatty acid (PUFA) supplementation in particular may have therapeutic potential for *H. pylori* treatment *via* multiple mechanisms, including suppression of inflammatory cytokines and enzymes and direct bacteriocidal effects *via* disruption of cell membranes^[16–18]^. 10-hydroxy-cis-12-octadecenoic acid (10-HOE; a linoleic acid derivative) and docosahexaenoic acid (DHA) have both been shown to inhibit *in vitro* growth of *H. pylori* and gastric infection in mouse models^[19, 20]^. Nevertheless, the effects of *H. pylori* infection on fatty acid metabolism in the gastrointestinal tract are not completely understood. Furthermore, the extent to which such metabolomic changes may relate to alterations in the gastrointestinal microbiota are unknown. Notably, short-chain fatty acids (SCFAs) produced by gut microbiota are thought to play a role in intestinal homeostasis and regulation of immune function, and dysbiosis in the setting of inflammatory bowel disease (IBD), which is associated with decreased levels of these important molecules^[21, 22]^. Exploration of changes in fatty acid production and metabolism associated with *H. pylori*-mediated alterations of the fecal microbiome is warranted, as it may reveal strategies to inhibit the cascade of events leading to neoplasia.

Previously, we studied the pattern of *H. pylori* prevalence and antibiotic resistance in our socioeconomically challenged and underprivileged inner-city community^[23]^. As a next step, we then sought to characterize the gut microbiome associated with *H. pylori* infection in our patients. We further explored interactions among *H. pylori* infection, the gut microbiota, and the gut metabolome by comparing stool fatty acid profiles between subjects with and without *H. pylori* infection. Our overarching goal is to identify potential targets for intervention that could improve *H. pylori* eradication rates and limit the consequences of repeated treatment failure.

## MATERIALS AND METHODS

### Patients and sample collection

Patients presenting to the Cooper Digestive Health Institute in Camden, New Jersey and diagnosed with *H. pylori* infection *via* biopsy were recruited for our study. *H. pylori* patients at least 18 years of age and diagnosed *via* biopsy between June 1, 2017 and December 31st, 2019, were eligible for inclusion. Patients less than 18 years old, pregnant women, patients taking antibiotics at the time of recruitment or sampling, and non-English speaking patients were excluded from participation. Control participants were recruited from a healthy population, also seen at Cooper Health Institute for well visits. Informed consent was obtained by one of several IRB-approved team members. Surveys about demographic information and relevant medical history were collected. Information on diagnosis, disease features, medication, history of antibiotic use, and treatment regimen were collected from the electronic medical record. Patients were asked to provide a stool sample before initiating eradication therapy, and samples were collected and stored in Stool Nucleic Acid Collection and Preservation Tubes (Cat. No. 45630, 45660, Norgen Biotek Corp., Thorold, Ontario, Canada) prior to freezing at –80 ^o^C as described previously^[23]^.

### 16S rRNA gene sequencing

Total genomic bacterial DNA extraction was performed from frozen fecal samples using the Qiagen DNeasy PowerSoil HTP Kit (Cat. No. 12955-4, Qiagen, Valencia, CA, USA). Adequate DNA yield was confirmed using NanoDrop spectrophotometry. For high throughput sequencing, 515F–806R Golay barcodes were used to target the hypervariable V4 region of the 16S rRNA gene, which is highly conserved and ideal for gut microbiome analysis. Marker genes in isolated DNA were PCR-amplified using GoTaq Master Mix (Cat. No. M5133, Promega, Madison, WI, USA) as described in Caporaso et al.^[24]^ PCR products were cleaned and normalized using the SequalPrep^TM^ Normalization Plate Kit (Cat. No. A1051001, ThermoFisher, Waltham, MA, USA) following manufacturer’s instructions. Purification of PCR product amplicons was performed with the QIAquick PCR Purification Kit (Cat. No. 28104, Qiagen). Quantification of the PCR amplified library was performed with the Invitrogen^TM^ Quant-iT^TM^ PicoGreen^TM^ dsDNA Assay Kit (Cat. No. P7589, ThermoFisher). 16S rRNA gene sequencing was performed on normalized pooled amplicons using the Illumina MiSeq System using a MiSeq reagent kit v2 (300 cycles; Cat. No. MS-102-2002, Illumina Inc.). FASTQ files for reads 1 (forward), 2 (reverse), and the index (barcode) read were generated using the BCL-to-FASTQ file converter bcl2fastq (ver. 2.17.1.14, Illumina, Inc.). Sequencing of the samples was conducted at the University of Colorado Boulder BioFrontiers Next-Gen Sequencing core facility.

### Sequencing data analysis

Analysis was performed using Python-based packages (Python 3.6.12), QIIME 2 2019.4 (denoising and taxonomy assignment), Quantitative Insights Into Microbial Ecology (QIIME) 2 2020.11 (statistical testing), and Linear Discriminant Analysis using LEfSe10^[25, 26]^. Sequences were de-multiplexed then filtered and clustered into sub-operational taxonomic units (sOTUs) using QIIME 2 DADA2^[27]^. A masked phylogenetic tree was created using MAFFT via QIIME2^[28, 29]^. Taxonomy was assigned using a naïve-Bayes classifier based on the latest Greengenes 16S rRNA gene database available as of August 2019 *via* the QIIME 2 interface (gg_13_8). Additional Python packages (SciPy.stats, Scikit-bio) were used for statistical tests on QIIME 2-generated data. Inter-class comparisons were made based on attributes of our population including: *H. pylori* status, gender, age group, race, education, and alcohol use. Alpha and beta diversity analyses were performed in QIIME 2, rarefied to an even sampling depth of 22,400^[30]^. Alpha diversity was assessed by Faith’s phylogenetic diversity, chao1 index, and observed operational taxonomic units (OTUs). Differences in alpha diversity between groups were determined by Kruskal-Wallis non-parametric rank test for categorical variables or by Fisher Z transformation on Spearman rho values for differences between alpha diversity correlations with numeric data across groups. Bacterial composition (beta diversity) was analyzed by principal coordinate analyses (PCoA) of both unweighted and weighted unique fraction metric (UniFrac) phylogenetic distance matrices^[31–34]^. Beta diversity was assessed for statistical significance between groups using Monte Carlo permutations [Adonis and permutational multivariate analysis of variance (PERMANOVA)], and dispersion of group samples was assessed using permutational analysis of multivariate dispersions (PERMDISP)^[35, 36]^. Comparisons of relative abundances of microbial taxa between groups were performed using Linear Discriminant Analysis (LDA) Effect Size (LEfSe) with the Huttenhower *et al.* online interface^[37]^. Taxa with an LDA value of 2.0 or greater and a two-tailed *p* value ≤ 0.05 with Kruskal-Wallis and pairwise Wilcoxon analyses were considered significantly enriched. The code is available at https://github.com/sterrettJD/H-pylori-microbiome-analysis.

### Non-targeted fatty acid analysis

Collected fecal samples were stored at –80 ^o^C until analysis. Sample preparation and analysis were performed using gas chromatography–mass spectrometry (GC-MS) by Metabolon, Inc. (Durham, NC, USA). This method (Metabolon TAM112-002) measures the total content of 30 fatty acids in stool samples after conversion into their corresponding fatty acid methyl esters (FAMEs). The measured concentrations are provided in weight corrected μg/ g of sample. NQ values were treated as a concentration of 0.001 μg/mL.

Briefly, stool samples were homogenized and the suspension was weighed out in a 100 mg aliquot in a test tube. Liquid/liquid extraction was performed to extract fatty acids and remove the nucleic acid preservative. A 250 μL aliquot of each extract was transferred to a clean analysis tube. The solvent was removed by evaporation under a stream of nitrogen. Internal standard solution was added to the dried sample extracts, quality controls (QCs), and calibration standards. The solvent was again removed by evaporation under a stream of nitrogen. The dried samples and QCs were subjected to methylation/transmethylation with methanol/sulfuric acid, resulting in the formation of the corresponding methyl esters (FAME) of free fatty acids and conjugated fatty acids. The reaction mixture was neutralized and extracted with hexanes. An aliquot of the hexanes layer was injected onto a 7890A/5975C GC/MS system (Agilent Technologies, Santa Clara, CA, USA). Mass spectrometric analysis was performed in the single ion monitoring (SIM) positive mode with electron ionization. Quantitation was performed using both linear and quadratic regression analysis generated from fortified calibration standards prepared immediately prior to each run. Raw data were collected and processed using Agilent MassHunter GC/MS Acquisition B.07.04.2260 and Agilent MassHunter Workstation Software Quantitative Analysis for GC/MS B.09.00/ Build 9.0.647.0. Data reduction was performed using Microsoft Office 365 ProPlus Excel. The metabolomic data set for 19 test and 12 control samples is included in the Supplementary materials (Full data set *H. pylori*). Volcano plot data are also included as Supplementary Tables S1, S2 and S3 (Table S1: Volcano plot data with all *Helicobacter* data set, Table S2: Volcano plot data with High Bacteroidetes and Table S3: Volcano plot data with High Firmicutes).

### Procrustes analysis

Multi-dimensional scaling (MDS, using the Python package Scikit-learn) was performed on Bray-Curtis distance matrices created from both the fecal metabolome and fecal microbiome datasets (using the Python package SciPy.distance), and a Procrustes test was performed on the metabolomics and microbiome MDS coordinates. As described by Peres-Neto and Jackson, a Procrustes randomization test (PROTEST) was performed by randomly sampling the MDS coordinates and performing a Procrustes test on the reordered coordinates 10,000 times, and the *p* value was calculated based on the portion of randomized Procrustes tests with resulting *m^2^* (Gower’s statistic) scores lower than the Procrustes *m^2^* score of the observed datasets^[38]^.

### Ethics approval

This study received approval by the Cooper Health System Institutional Review Board (IRB) (17-077EX) and all the steps were carried out as per the standards set by the IRB.

## RESULTS

### Fecal microbiome analysis

Selected demographic attributes of our participants are presented in Table 1. Fecal microbiome analysis of 35 stool samples collected from 19 *H. pylori* patients and 16 healthy control subjects was carried out. The mean number of reads per sample was 269,462 with a minimum of 22,483 reads/sample. Rarefaction was set to 22,400 for even sampling based on this minimum. We analyzed the effect of each of the factors included in Table 1 on alpha and beta diversity. Age was the only factor that showed statically significant differences (p value of ≤ 0.05) in the *H. pylori* patients compared to the control subjects.

**Table 1.** Patient Demographics and Clinical Characteristics.

|  | Total (*n* = 35) | *H. pylori* patients (*n* = 19) | Control (*n* = 16) |
| --- | --- | --- | --- |
| Sex |  |  |  |
| Male | 19 (54%) | 13 (68%) | 6 (38%) |
| Female | 16 (46%) | 6 (32%) | 10 (62%) |
| Age |  |  |  |
| Ages 18-40 | 15 (43%) | 3 (16%) | 12 (75%) |
| Age >40 | 20 (57%) | 16 (84%) | 4 (25%) |
| Highest education |  |  |  |
| Less than high school | 2 (6%) | 2 (11%) | 0 (0%) |
| High school | 12 (34%) | 11 (58%) | 1 (6%) |
| Undergraduate | 4 (11%) | 2 (11%) | 2 (13%) |
| Graduate | 15 (43%) | 3 (16%) | 12 (75%) |
| Post-graduate | 1 (3%) | 0 (0%) | 1 (6%) |
| Race |  |  |  |
| African American | 8 (23%) | 6 (32%) | 2 (12%) |
| Asian American | 2 (6%) | 0 (0%) | 2 (12%) |
| Caucasian | 13 (37%) | 1 (5%) | 12 (75%) |
| Non-White Hispanic | 8 (23%) | 8 (42%) | 0 (0%) |
| Other | 3 (9%) | 3 (16%) | 0 (0%) |
| Smoker | 5 (14%) | 4 (21%) | 1 (6%) |
| Past *H. pylori* treatment | 1 (3%) | 1 (5%) | 0 (0%) |
| Duodenal ulcer | 1 (3%) | 1 (5%) | 0 (0%) |
| Non-ulcer dyspepsia | 14 (40%) | 10 (53%) | 4 (25%) |

### Analysis of alpha diversity

Chao1 index alpha diversity between *H. pylori* patients and control subjects and regression analysis of observed OTUs by age are shown in Figure 1. Alpha diversity, which estimates the diversity of a microbial community within a sample, incorporates the number of different OTUs and their respective abundance in each sample. Alpha diversity was lowest in *H. pylori* patients over 40 years of age, who had significantly lower Chao1 scores compared to control subjects over 40 years of age (Kruskal-Wallis *H* = 6.036, *p* = 0.014). *H. pylori* patients over 40 years old also had lower Chao1 scores approaching statistical significance when compared to *H. pylori* patients under 40 (Kruskal-Wallis *H* = 2.813, *p* = 0.094). Additionally, the correlation of age with observed OTUs (Figure 1B), Faith’s phylogenetic diversity, and Chao1 was consistently different between *H. pylori* patients *versus* control subjects, as evidenced by a Fisher *Z* transformation on Spearman rank correlation *rho* values (observed OTUs: *H. pylori rho* = −0.21 vs control *rho* = 0.50, Fisher *p* = 0.041; Faith: *H. pylori rho* = −0.04 vs control *rho* = 0.57, Fisher *p* = 0.067; Chao1: *H. pylori rho* = −0.15 vs control *rho* = 0.44, Fisher *p* = 0.097).

**Figure 1.**
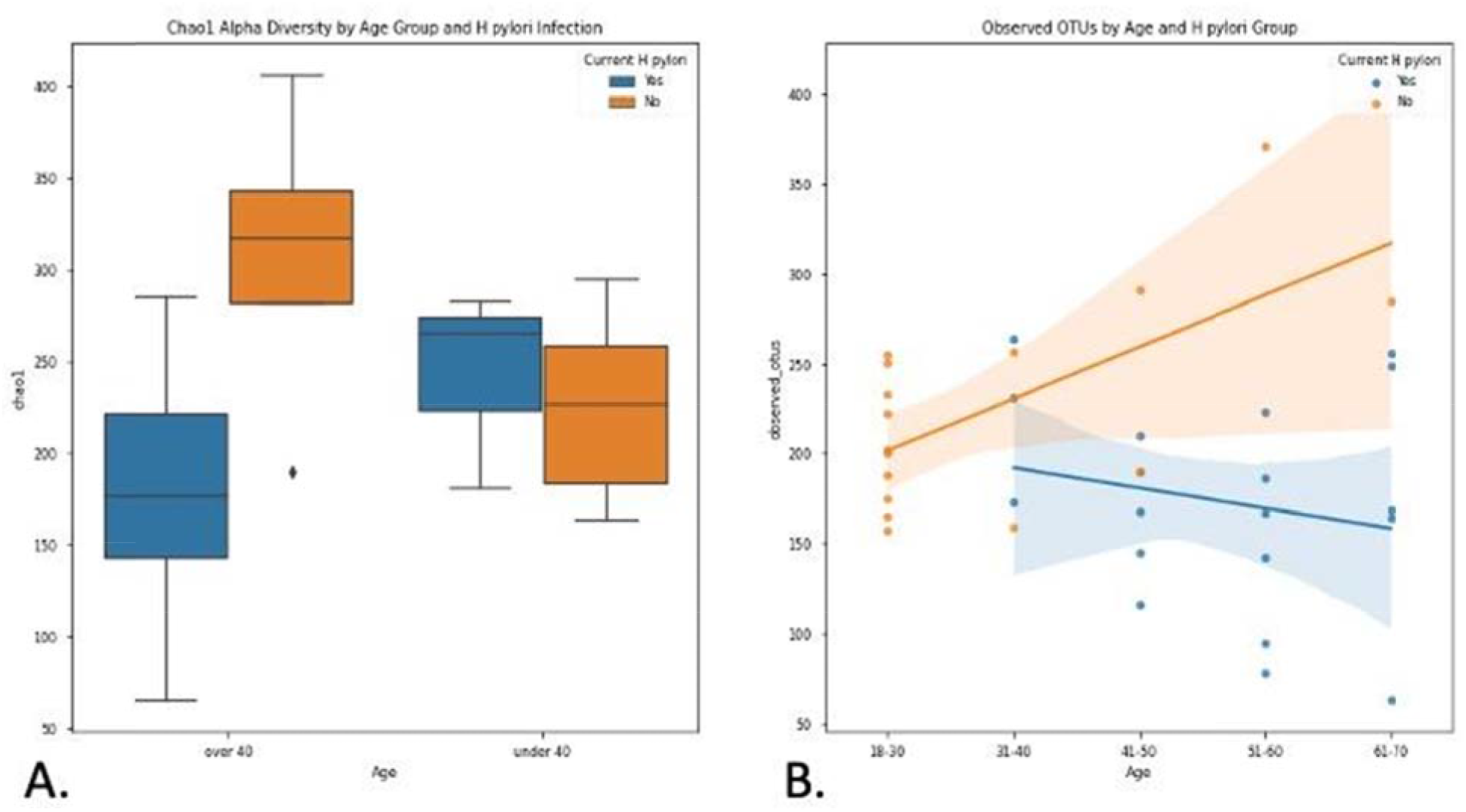
Chao1 index alpha diversity and observed OTUs by *H. pylori* status stratified by age. **A**: Chao1 index alpha diversity is compared between *H. pylori* patients (blue) and control subjects (orange), subgrouped by age. *H. pylori* patients over 40 years old (n=16) had significantly lower Chao1 scores compared to control subjects over 40 (n=4) (Kruskal-Wallis *H* = 6.036, *p* = 0.014). Boxes represent 1st and 3rd quartiles, and central lines represent median values. Whiskers represent non-outlier high and low values. Points show outliers, which were determined by having a distance from the 1st or 3rd quartile greater than 1.5 times the interquartile range. B: Observed OTUs (operational taxonomic units) are compared between *H. pylori* patients (blue, n=19) and control subjects (orange, n=16), as a function of age. Shading around the regression lines indicates 95% confidence intervals.

Next we analyzed beta diversity of our samples, which estimates how samples differ from each other. An unweighted UniFrac PCoA plot of *H. pylori* patients *versus* control subjects is shown in Figure 2. Adonis testing revealed that current *H. pylori* infection represented 7% of the variation in unweighted UniFrac beta diversity *(F* = 2.523, *R^2^* = 0.071, *p* = 0.002). Additionally, unweighted UniFrac PERMANOVAs identified significant differences between *H. pylori* patients and control subjects *(F* = 2.523, *p* = 0.001). Furthermore, unweighted UniFrac PERMDISP did not reveal differences in dispersion among *H. pylori* patients *versus* control subjects (*F* = 0.0506, *p* = 0.809), but weighted UniFrac PERMDISP did (*F* = 6.82955, *p* = 0.017). We also explored the effect of the use of proton pump inhibitors (PPIs) *via* beta diversity analysis as PPIs are the medications commonly given to *H. pylori* patients and we had previously noted their influence on the antibiotic resistance of *H. pylori*^[23]^. Unweighted UniFrac PERMANOVA identified differences between PPI users and non-users across all participants (*F* = 1.939, *p* = 0.012), though the magnitude of the effect was smaller than that seen for *H. pylori* infection as described above. It should also be noted that only four *H. pylori* patients and one control subject were on PPI at the time of study and sample collection, so its contribution to the results described here may not be significant.

**Figure 2.**
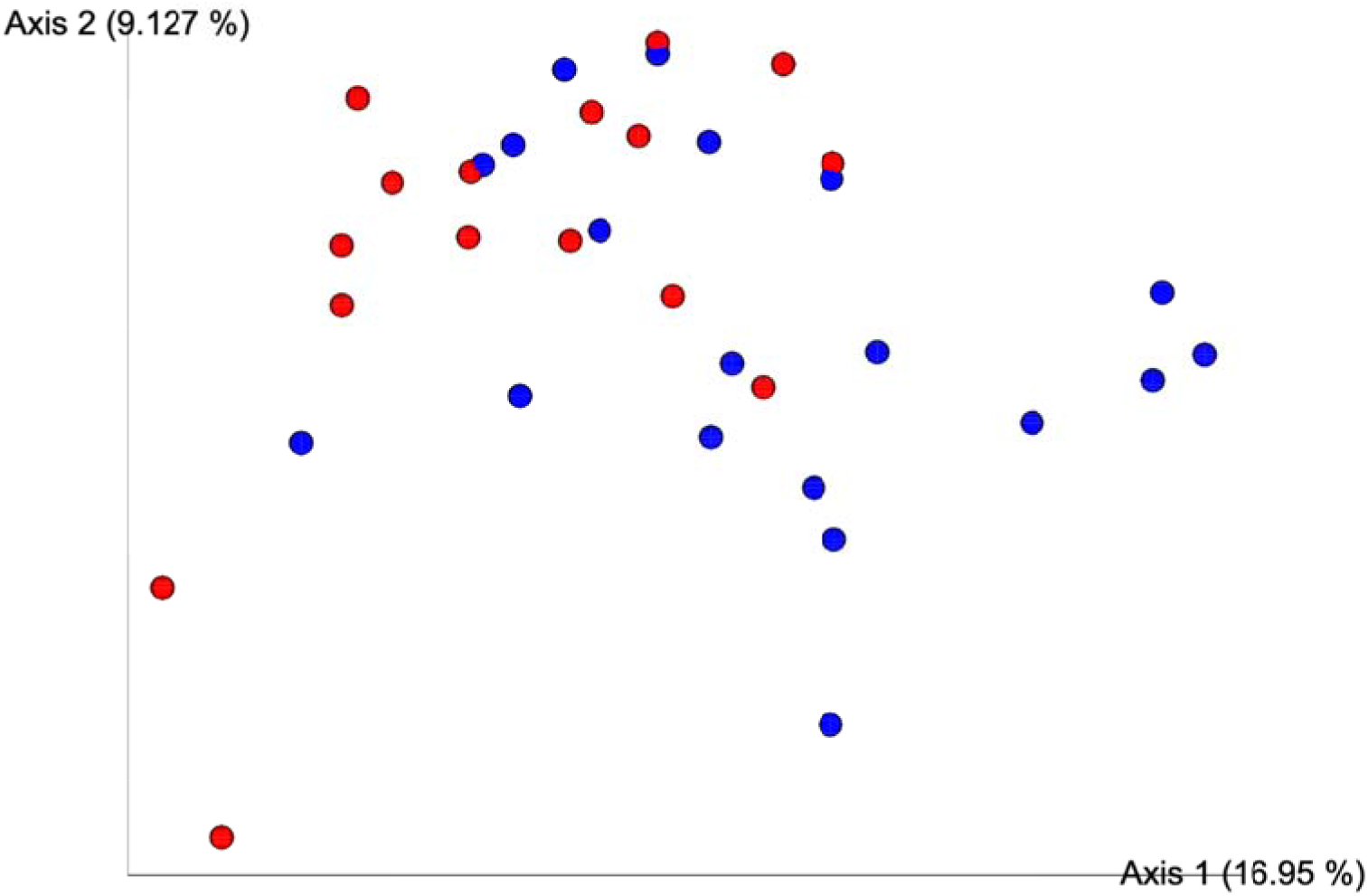
Beta diversity analysis by unweighted UniFrac PCoA. Unweighted Principal Coordinate Analysis (PCoA) plot is shown comparing *H. pylori* patients (blue, n=19) to control subjects (red, n=16). Significant differences between *H. pylori* patients and control subjects were identified by unweighted UniFrac PERMANOVA (*F* = 2.523, *p* = 0.001). The proportion of variance explained by each principal coordinate axis is denoted in the corresponding axis label; PCo1 explains 16.95% of the variability and PCo2 explains 9.13% of the variability.

### Linear discriminant analysis effect size

LEfSe scores for taxa that were differentially distributed across *H. pylori* patients *versus* control subjects are shown in Figure 3. Thirty-eight taxa had lower relative abundance in *H. pylori* patients relative to healthy control subjects, and six taxa had higher relative abundance in *H. pylori* patients relative to healthy control subjects. Taxa that were enriched in *H. pylori* patients include *Atopobium*, Gemellaceae, Micrococcaceae, Gemellales and *Rothia* (*R. mucilaginosa*). Figure 4 shows the arcsin square root transformed relative abundances of each phylum, grouped by *H. pylori* status. Notably, relative abundance of the phylum Verrucomicrobia was decreased in *H. pylori* patients compared to control subjects.

**Figure 3.**
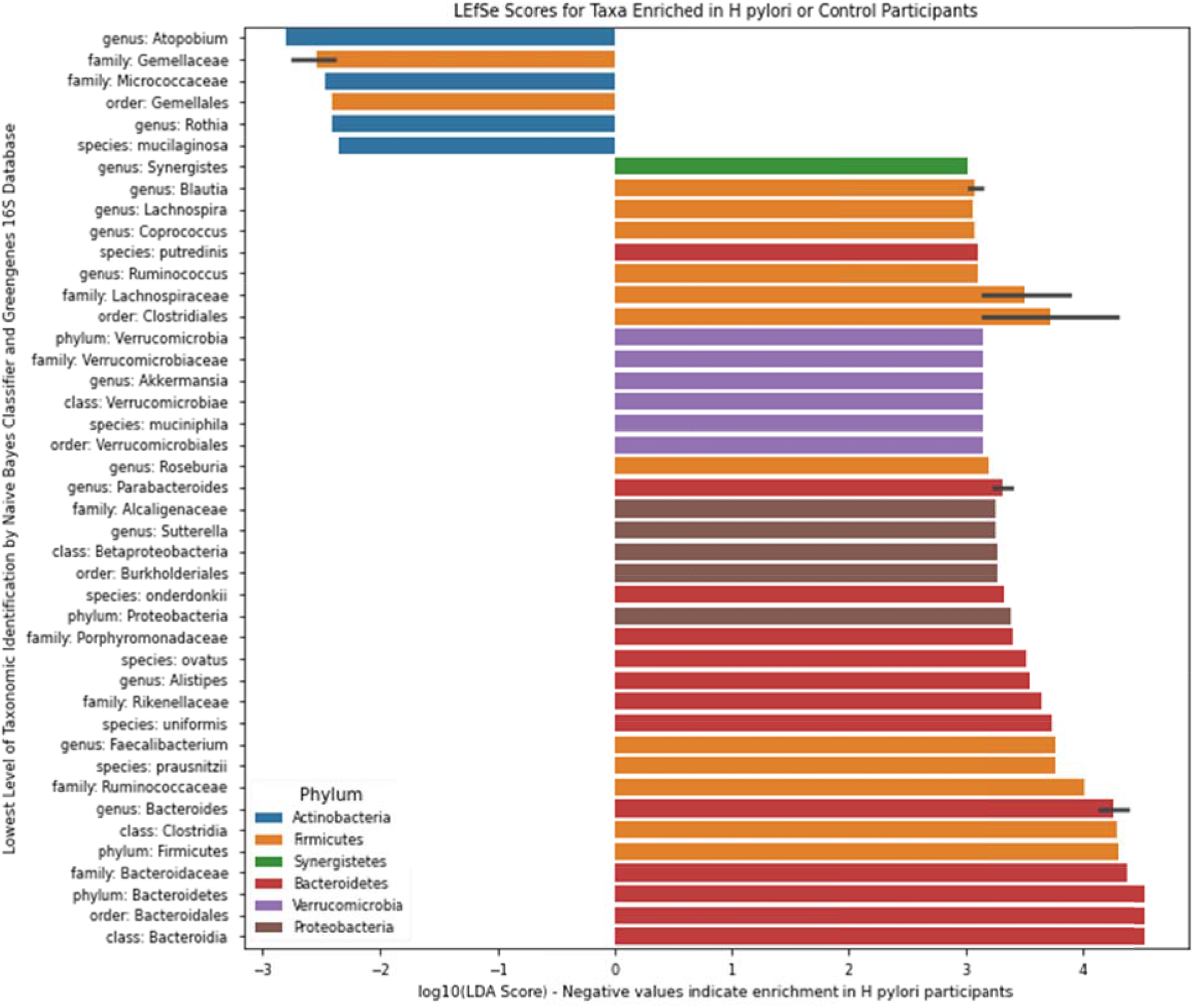
Linear discriminant analysis effect size (LEfSe). LEfSe scores for taxa that were differentially distributed across *H. pylori* patients (n=19) *versus* control subjects (n=16) *via* Kruskal-Wallis and Wilcoxon tests with two-tailed α = 0.05. Negative values represent taxa that were enriched in *H. pylori* patients, whereas positive values represent taxa that were enriched in control subjects. Error bars represent a 95% confidence interval and are only present if the listed taxonomic identity was the lowest level classified for multiple OTUs highlighted by LEfSe. Color represents the phylum.

**Figure 4.**
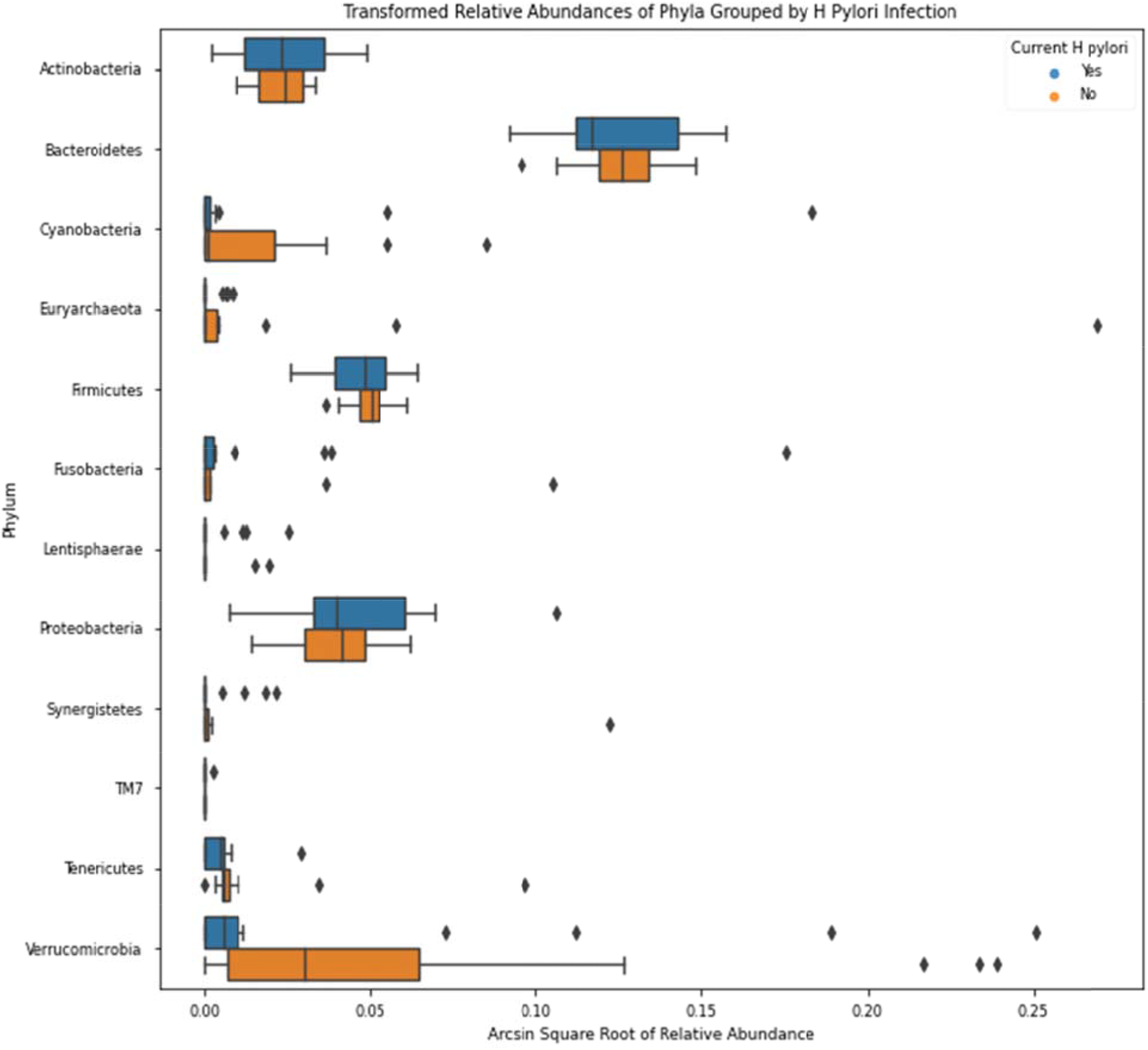
Relative phylogenic abundance. Arcsin square root transformed relative abundances of each phylum, grouped by *H. pylori* status. Bacterial relative abundances of *H. pylori* patients (n=19) are shown in blue, whereas healthy controls (n=16) are shown in orange. Boxes represent the 1st and 3rd quartiles, and vertical lines in the middle of the boxes represent median values. Whiskers represent the lowest and highest non-outlier values. Points show outliers, which were determined by having a distance from the 1st or 3rd quartile greater than 1.5 times the interquartile range. Of note, relative abundances of Verrucomicrobia were significantly lower in *H. pylori* patients compared to healthy controls (Kruskal-Wallis H = 4.455, p = 0.034).

### Procrustes analysis

The Procrustes randomization test was carried out to investigate the relationship between the fecal microbiome and fecal metabolome. Procrustes analysis identified a significant relationship between the two datasets, as the resulting *p*-value of 0.017 indicates that the correspondence of each participant’s fecal microbiome to their fecal metabolome was better than 98% of randomly sampled simulations (Figure 5).

**Figure 5.**
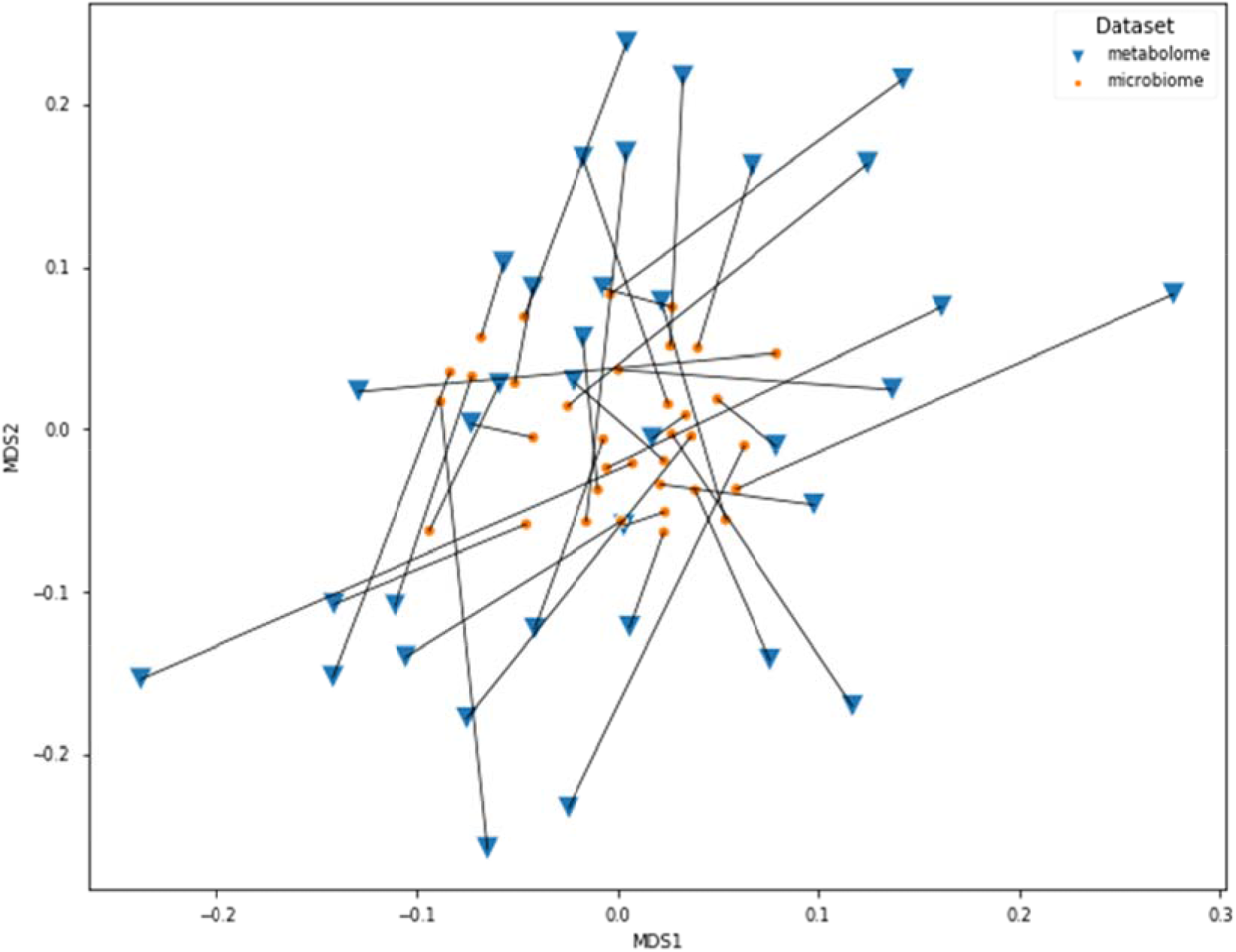
Procrustes analysis. Connections between the fecal microbiome and fecal metabolome Bray-Curtis MDS plots. The microbiome data shown here have been transformed by the Procrustes analysis to minimize disparity between the two datasets, and lines have been drawn to indicate microbiome and metabolome from the same participants. Orange dots represent the microbiomes, whereas blue triangles represent the metabolomes. The Procrustes randomization test showed a significant relationship between the datasets (*m^2^* = 0.86, *p* = 0.017).

### Fecal Fatty acid analysis

To evaluate the impact of *H. pylori* infection on fatty acid composition, we performed non-targeted metabolomics analysis of a panel of 30 fatty acids including long chain fatty acids (LCFAs), monounsaturated fatty acids (MUFAs) and polyunsaturated fatty acids (PUFAs). We prepared a volcano plot evaluating changes in fatty acid profile for the entire population of *H. pylori* patient samples relative to healthy control subjects (Figure 6a). In addition, we prepared separate volcano plots for subgroups of the *H. pylori* patients with high Bacteroidetes and high Firmicutes based on 16S rRNA gene analysis (Figure 6b and c). See Supplementary materials (Tables S1, S2, S3) for the full volcano plot data. Volcano plots allow for the plotting of *p*-values *versus* fatty acid fold change from control samples for all evaluated fatty acids.

**Figure 6.**
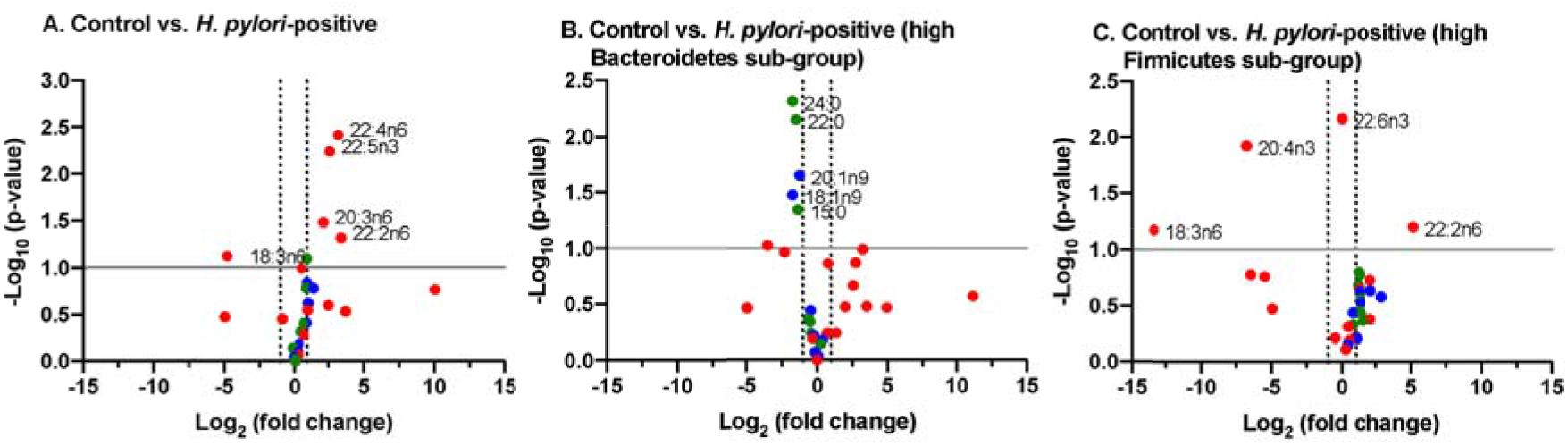
Volcano plots of composition of evaluated fatty acids. A. Control vs. entire *H. pylori* patient group. B. Control *versus* sub-group of *H. pylori* patients identified as having high ratio of Bacteroidetes to Firmicutes (*n* = 7 of 19 patients). C. Control *versus* sub-group of *H. pylori* patients identified as having high ratio of Firmicutes to Bacteroidetes (*n* = 4 of 19 patients).

Stool samples from *H. pylori* patients showed increases in several fatty acids including the PUFAs 22:4n6, 22:5n3, 20:3n6 and 22:2n6 while decreases were noted in fatty acids including the PUFA 18:3n6, relative to healthy control subjects. The pattern of changes in fatty acid concentration was correlated to the Bacteroidetes:Firmicutes ratio (determined by 16S rRNA gene analysis). Among the sub-population identified to have high Bacteroidetes (Bacteroidetes:Firmicutes ratio ≥ 1 standard deviation above control ratio, *n*=7) we found several fatty acids that were diminished from control including the LCFAs 24:0, 22:0 and 15:0 along with MUFAs 20:1n9 and 18:1n9. By contrast, in the subpopulation identified to have high Firmicutes (Firmicutes:Bacteroidetes ratio ≥ 1 standard deviation above control ratio; *n*=4) a distinct set of fatty acids were found to be significantly increased and decreased from controls. In this subpopulation we observed increases in the PUFA 22:2n6 and decreases in PUFAs 18:3n6 and 20:4n3.

## DISCUSSION

Our sample size is modest due to the stringent limitations posed by the socioeconomic and cultural attributes of our patient population and logistic challenges. Camden County has the lowest median income and highest poverty and unemployment rates in southern New Jersey. It also has the lowest educational attainment and the greatest socioeconomic disparities among ethnic minority groups. More than one third of the population in Camden City, where our hospital is located, lives below the national poverty line, and the median household income is USD $26,105^[39]^. Our local community is thus considered one of the poorest and most economically distressed communities in the United States. A significant fraction of adults over 25 years old (almost one quarter) have not completed high school. It is possible that this influences patients’ appreciation of the relevance of the study and willingness to participate. Please note that in the present study, 13 out of 19 participating patients had education level of high school or less. A significant portion of potential participants were excluded due to non-English-speaking status, thus being unable to partake in the informed consent process. Logistic challenges imposed by the process of stool sample collection also limited the number of subjects. A large number of potential participants were unable to return to the clinic for stool sample delivery due to poor transportation access or inability to leave employment or domestic responsibilities during the window for sample collection (after diagnosis but prior to initiation of antibiotics). Nonetheless, the present study yielded meaningful results.

*H. pylori* infection was associated with significantly decreased fecal microbial diversity among patients over the age of 40 years, while the control subjects had the highest microbial diversity. Correlations between age and multiple metrics of microbial alpha diversity were consistently different between *H. pylori* patients and control subjects. *H. pylori* was also shown to account for a significant portion of the variation in beta diversity between infected patients and control subjects. Collectively, these results could suggest an *H. pylori*-mediated transformation of the fecal microbiome that progresses with age and becomes apparent by mid-adulthood. Central to this transformation is a decline in fecal microbial diversity that may be the result of longstanding infection in which *H. pylori* alters the gastrointestinal environment at the expense of other taxa. This conclusion is consistent with those of other studies that attributed decreased diversity of gastric microbiota to the ability of *H. pylori* to outcompete other species^[9, 14, 40]^. Secretion of antibacterial peptides and local pH alterations have been proposed as mechanisms that may account for the alteration of the gastric microbiota diversity and community structure by *H. pylori*^[41]^. The impact of *H. pylori* on fecal microbial diversity observed in our study may be reversible, just as a recent study showed that *H. pylori* eradication could restore gastric microbial diversity to levels seen in uninfected controls^[42]^.

Conversely, the data may reflect a preceding decrease in gut microbial diversity that facilitates *H. pylori* infection, possibly mediated by other factors or exposures that accumulate with age. In other words, patients who experience decreased gut microbial diversity as they age, potentially mediated by diet or chronic inflammation, may be more prone to *H. pylori* infection, particularly by mid-adulthood. In our previous study of 2,014 *H pylori* patients, we observed that 80% of our patients were above the age of 40 years^[23]^. Antibiotic exposure is known to alter the gut microbiome, and increased rates of antibiotic resistant-*H. pylori* among older adults have been attributed to increased antibiotic use compared to younger patients^[2, 43, 44]^. Conceivably, decreased gut microbial diversity induced by cumulative antibiotic exposure could create an intragastric niche for *H. pylori*, and antibiotic resistant *H. pylori* strains in particular, to thrive.

LEfSe showed that multiple taxa were differentially enriched or depleted in stool from *H. pylori* patients compared to controls. The significance of differential expression of these taxa is unknown; however, certain taxa stand out. The enrichment of *Atopobium* is notable in the context of a metagenomic analysis which revealed increased *Atopobium parvulum* abundance among patients with colorectal intramucosal carcinomas^[45]^. Although *H. pylori* is predominantly associated with gastric cancers, a recent metaanalysis suggested a significantly increased risk of colorectal cancer as well, and our data suggest that *H. pylori*-associated increases in *Atopobium* abundance may have a role in the distal gut carcinogenesis^[46]^.

Over-representation of *Rothia mucilaginosa*, a commensal oral microbe, has previously been observed in fecal samples from patients with primary sclerosing cholangitis (PSC)^[47]^. Because of the sensitivity of *R. mucilaginosa* to gastric fluid, Bajer *et al*. hypothesized that its presence in PSC patients represented contamination of the intestine *via* previous endoscopic retrograde cholangiopancreatography^[47]^. The enrichment of *Rothia* sp. in fecal samples from our *H. pylori* patients may reflect the potent gastric acid suppression by *H. pylori*, allowing translocation of swallowed oral flora to the distal gut. The clinical consequences of the colonization of the distal gut by *Rothia* sp. are unknown, although the bacteria has been implicated as a cause of myriad syndromes, including dental caries, pneumonia, and bacteremia, particularly in immunocompromised patients^[48]^. The increased abundance of Gemellaceae in *H. pylori* patients is consistent with the observation of increased organisms from the *Gemella* genus in patients with current *H. pylori* infection reported by Gao et al^[49]^. Microbial shift that included the expansion of Gemellaceae in the Crohn ileum has been reported by other groups^[50, 51]^.

Interestingly, the phylum Verrucomicrobia was significantly depleted in *H. pylori* patients. Verrucomicrobia includes *Akkermansia muciniphila*, a mucus-residing commensal bacterium of the large intestine. *A. muciniphila* is an obligate chemoorganotroph, utilizing mucus as a sole carbon, nitrogen, and energy source that produces SCFAs including acetate, propionate, and, to a smaller extent, 1,2-propanediol^[52, 53]^. Decreased relative abundance of Verrucomicrobia may suggest disruption of the gut mucosal environment by *H. pylori*. In a mouse stress model, exposure to chronic psychosocial stress resulted in expansion of *Helicobacter* spp., which has been demonstrated to proliferate in response to glucocorticoid administration as well as psychosocial stress exposure^[54]^. Specifically, stress-induced increases in *Helicobacter* spp. were associated with increases in other *Proteobacteria* spp., including unidentified genera belonging to the Enterobacteraceae and Desulfovibrionaceae families. Concurrently, relative abundance of *Mucispirillum*, an obligate mucus-residing bacterium, decreased; a decline of *Mucispirillum* in rodents is associated with early disruption of the gastrointestinal surface mucus layer and a prolonged delay to recovery after the period of pathogen clearance^[55]^. Evidence of disruption of the gut mucosa by *H. pylori* infection is also consistent with decreased relative abundance of *Faecalibacterium prausnitzii*. *F. prausnitzii* is an important gram-positive human commensal that produces butyrate and other SCFA through the fermentation of dietary fiber. Reduced relative abundance of *F. prausnitzii* has been associated with inflammatory conditions including Crohn’s disease, obesity, asthma, and stress-related psychiatric disorders in which inflammation is a risk factor, such as major depressive disorder^[56–59]^.

The gastric microbiome differs significantly from those of other sites along the gastrointestinal tract^[60]^. Nevertheless, *H. pylori* infection is thought to indirectly alter distal gut microbial community structure^[61]^. Recent studies have identified significant differences in bacterial communities obtained from human duodenal and stool samples associated with *H. pylori* infection^[62, 63]^. Schulz *et al.* identified a few phylotypes that were detected at significantly different rates in duodenal aspirates or biopsies of *H. pylori*-positive individuals compared to controls, and this finding may be related to *H. pylori’s* association with duodenal ulcers^[62]^. In contrast to our study, Dash *et al.* observed increased alpha diversity among stool samples from *H. pylori*-positive individuals, although they found no effect on beta diversity^[63]^. Finally, Frost *et al.* found significant changes in fecal microbiome diversity and composition as measured by alpha and beta diversity, respectively, in *H. pylori* positive individuals compared to controls, and the latter association was strongly correlated with *H. pylori* stool antigen load^[64]^. Differences in the study populations, including geographic, cultural, and dietary differences, may account for these conflicting findings regarding *H. pylori’s* effects on the distal gut microbiome, as these factors are known to significantly impact fecal microbiome composition^[65]^. Moreover, methodological differences in analyses of alpha and beta diversity and relative taxonomic abundance further inhibit comparison of study results. Nevertheless, our results add to the growing evidence suggesting that *H. pylori* is associated with alterations in distal gut microbiota, particularly in those over 40 years of age.

Strategies to promote increased gut microbial diversity among patients with *H. pylori* infection, particularly those over 40 years of age, may hold therapeutic potential in eradicating infection, and some clinical trials with adjunctive probiotic formulas have shown promising results in this regard^[66]^. Likewise, treatment of gut dysbiosis in patients with *H. pylori* infection may limit its associated pathologies. Reduced fecal microbiome diversity is associated with multiple disease markers including increased adiposity, insulin resistance, dyslipidemia, and a pro-inflammatory state ^[67]^. *H. pylori* infection is associated with higher rates of diabetes, specifically, and addressing gut dysbiosis in such patients could conceivably aid primary dietary and pharmacologic strategies to achieve glycemic control and insulin sensitivity^[12]^. Rehabilitation of the gut microbiota in patients with *H. pylori* may also have a role in minimizing the risk of malignancy. Mouse studies suggest that the carcinogenic effects of *H. pylori* partly depend on interactions with the gut microbiome, and persistent infection with *H. pylori* may create a niche favorable for taxa that are found in increased abundance in gastric cancer, including *Lactobacillus* and *Lachnospiraceae*^[9, 68, 69]^. It has been suggested that lactic acid bacteria can influence gastric cancer by a number of mechanisms such as (i) supply of exogenous lactate, that acts as a fuel source for cancer cells, (ii) production of reactive oxygen species and N-nitroso compounds and (iii) by allowing colonization of carcinogenic non-*H. pylori* bacteria^[70]^. Accordingly, reversal of abnormalities in gut microbiota associated with *H. pylori* may also limit its carcinogenic potential. Finally, consideration must be given to the effects of *H. pylori* treatment on the gut microbiome, as data suggest that current therapies may be associated with dysbiosis and subsequent adverse effects^[71]^.

While evaluating changes in the long-chain fatty acids (LCFAs) palmitic (16:0) and stearic (18:0) concentrations, we found that palmitic acid concentrations were increased in our test samples compared to control samples (625.23±598.82 μg/g (*n*=18) *versus* 334.10±284.58 μg/g (*n*=12)), whereas concentrations of stearic acid were comparable between populations. Interestingly, when we evaluated the high Firmicutes (>1.4:1 Firmicutes: Bacteroidetes, *n*=4) sub-group compared to the high Bacteroidetes (>2.7:1 Bacteroidetes: Firmicutes*, n*=7) sub-group of our test population we found the concentrations of both these LCFAs to be diminished in the high Bacteroidetes population while the concentrations of both were enhanced in the high Firmicutes subpopulation. This is consistent with previous studies in mice where administration of LCFA-rich diets resulted in an observed increase in Firmicutes to Bacteroidetes ratio^[72]^. Ktsoyan *et al*. implicated a characteristic profile of serum LCFA concentrations in association with *H. pylori* infection compared to healthy controls^[73]^. It is plausible that diet can modulate the impact of *H. pylori* on the gut microbiome, which in turn may relate to different disease phenotypes, *i.e.*, asymptomatic infection, peptic ulcer disease, and gastric cancer.

The accumulation of the saturated LCFAs is regulated under normal physiological conditions by desaturation of 16:0 to 16:1n7, elongation of 16:0 to 18:0 and/or desaturation of 18:0 to 18:1n9 which can be further elongated to 20:1n9 (Figure 7). While only small changes in concentrations of these MUFAs were noted between our *H. pylori* patients and control subjects, the changes in MUFA concentrations in our high Firmicutes and high Bacteroidetes sub-populations showed more significant and consistent changes across the series of MUFAs. The concentration of 16:1n7, 18:1n9 and 20:1n9 were decreased in the high Bacteroidetes sub-group, whereas these MUFAs were slightly increased in the high Firmicutes sub-group. Interestingly, this contrasts published data on the benefits of high MUFA diets in animal models and humans. In previous studies, high MUFA diets have been reported to lead to increased ratios of Bacteroidetes to Firmicutes and were correlated with reductions in several disease indicators^[72]^. Together, these observations may suggest that either the patients in our high Bacteroidetes sub-population have severely limited MUFA production/intake, contrasting previous work on MUFA rich diets or that the endogenous biosynthetic pathways for MUFA production are dysregulated in this patient population.

**Figure 7.**
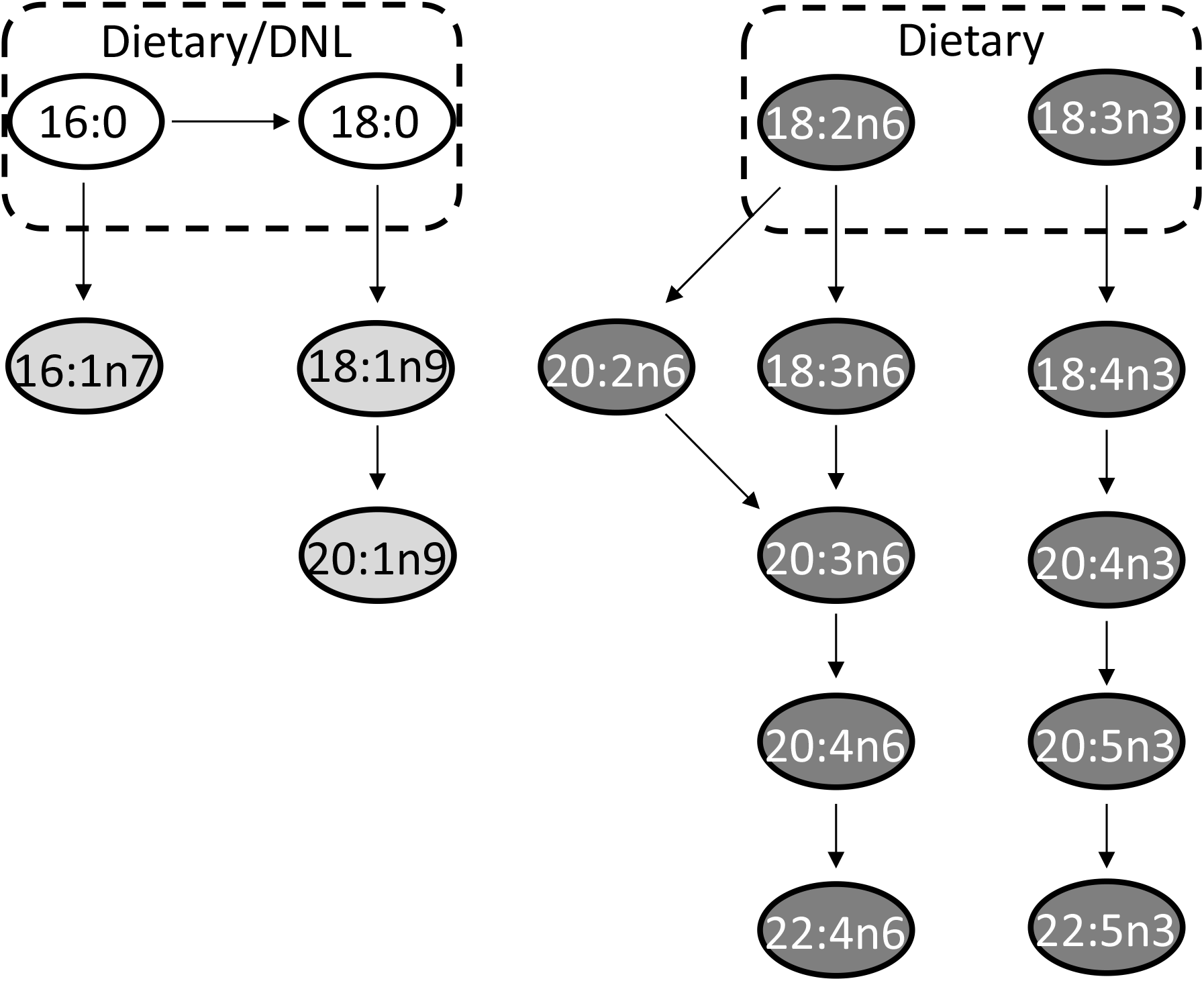
Major metabolic pathways. Selected long-chain fatty acids (LCFAs, no fill), monounsaturated fatty acids (MUFAs, light grey fill) and polyunsaturated fatty acids (PUFAs, dark grey fill) are highlighted. Although many of the fatty acids shown may be supplemented by diet, key parent members of these pathways acquired from *de novo* lipogenesis (DNL) and/or diet are indicated.

The range of PUFAs observed endogenously have essential physiological roles and are produced from dietary alpha-linolenic acid (18:3n3) and linoleic acid (18:2n6). We observed that both of these dietary PUFAs were increased in our test samples compared to the control samples. In humans, C18:2n6 is converted through multiple steps to eicosapentaenoic acid (EPA, 20:5n3) and docosahexaenoic acid (DHA, 22:6n3), whereas 18:3n3 proceeds to produce arachidonic acid (AA, 20:4n6) (Figure 7). As these two biosynthetic pathways share enzymes for their respective dehydration and elongation steps, they are antagonistic to each other. In general, a high 18:3n3 (n-3 fatty acid) diet is correlated with positive physiological effects whereas a high 18:2n6 (n-6 fatty acid) diet is correlated with metabolic dysbiosis and negative physiological effects^[74, 75]^. Accordingly a balanced dietary intake of n-6/n-3 fatty acids is recommended to be 1:1-2:1, although the average ratio in a typical Western diet has been reported to be >15:1^[76]^. In accordance with published ratios for n-6/n-3 fatty acid intake in Western diets, we find the ratio of 18:2n6/18:3n3 to be 22:1 in our control samples and 15:1 in our *H. pylori* patient test samples.

Endogenously, the essential dietary fatty acid 18:3n3 undergoes multiple dehydration and elongation steps to produce docosapentaenoic acid (DPA, 22:5n3) and docosahexaenoic acid (DHA, 22:6n3). For all intermediates along this biosynthetic pathway, with exception to 18:3n3 and 22:5n3, we observe concentrations at or below the limit of quantification for the majority of samples making an evaluation of the individual steps of this biosynthesis pathway difficult.

Contrasting the low concentrations observed for 18:3n3 and fatty acids produced along this biosynthetic pathway, higher overall concentrations were observed for fatty acids derived from 18:2n6, perhaps reflecting the higher concentration of dietary n-6 fatty acids as described earlier. In humans, the dietary fatty acid, 18:2n6 undergoes dehydration to produce 18:3n6. This intermediate undergoes a subsequent elongation to produce 20:3n6, which is dehydrated to 20:4n6 (arachidonic acid). Alternatively, 18:2n6 can undergo elongation first to produce 20:2n6, which is dehydrated to produce 20:3n6 (Figure 7).

We found that the early steps in the metabolism of 18:2n6 were significantly impacted in all of our *H. pylori* samples by comparison of downstream fatty acid metabolites to controls. Most notably, in all *H. pylori* test samples the concentration of 18:3n6 was significantly diminished. Concentration of 18:3n6 was 10.7±18.2 μg/ g (*n*=12) in the control samples and 0.36±1.50 μg/g (*n*=18) in the test samples. While we would anticipate that this deficiency in dehydration of 18:2n6 would lead to lower levels of fatty acid metabolites downstream of 18:3n6, this is not the case. In fact, by the following biosynthetic step, *i.e.*, elongation to produce 20:3n6, the fatty acid concentration of 20:3n6 in test samples was found to be higher than in controls. These higher concentrations were also reflected in arachidonic acid (20:4n6) and adrenic acid (22:4n6). It is plausible that the alternative *δ-8-desaturase* mediated route *via* 20:2n6 may be prominent in these *H. pylori* patient samples, which are marked by a deficiency in δ*-6-desaturase* mediated pathway *via* 18:3n6^[77]^.

As the burden of antibiotic resistance grows, populations like ours are impacted greatly by the high prevalence of *H. pylori* colonization. As eradication by a one-fits-all approach becomes more and more difficult, it becomes even more important to look towards strategies that can provide insight into community attributes that can help guide targeted therapies, predict likelihood of treatment outcomes, and possibly create strategies towards mitigating the impact of increased antibiotic usage. A greater, more individualized understanding of gut microbiome features among patients experiencing *H. pylori* colonization, especially in our underserved population, will pave the way for better community impact and reduce healthcare burdens of repeated treatment and mounting resistance. As gut dysbiosis is inextricably tied not only to *H. pylori* infection but also to the use of antibiotics itself, the cycle of treatment and re-treatment may very well only further exacerbate vulnerability to enteric pathogens. Strategies to better understand gut dysbiosis features among affected groups may create opportunities for targeted therapy, as well as a potential for creating predictive models for antibiotic selection, and even exploring restorative probiotic therapy to help lessen the adverse impact of eradication therapy.

## ACKNOWLEDGEMENT

This work was supported by the Camden Health Research Initiative grant to Sangita Phadtare. This grant is for academic research. Purchase of laboratory supplies, sequencing *etc* was carried out from this grant. The grant agency had no role in the design and execution of the study or in the interpretation of results. The grant agency did not have access to any data from the study. No other authors had grant support for research. We thank Drs. Robert Cooper, Anjali Mone and Sanket Patel for their help in the collection of some of the samples for the study; these were also approved in the IRB protocol.

## Abbreviations

10-HOE: 10-hydroxy-cis-12-octadecenoic acid
DHA: docosahexaenoic acid, 22:6 (n-3)
DNL: *de novo* lipogenesis
FAME: fatty acid methyl ester
GC-MS: gas chromatography–mass spectrometry
IBD: inflammatory bowel disease
IRB: Institutional Review Board
LDA: Linear Discriminant Analysis
LEfSe: Linear Discriminant Analysis (LDA) Effect Size (LEfSe)
MDS: multidimensional scaling
MUFA: monounsaturated fatty acid
OTU: operational taxonomic unit
PCoA: principal coordinate analysis
PERMANOVA: permutational multivariate analysis of variance
PERMDISP: permutational analysis of multivariate dispersions
PPI: proton pump inhibitor
PROTEST: Procrustes randomization test
PUFA: polyunsaturated fatty acid
QC: quality control
QIIME: Quantitative Insights Into Microbial Ecology
SCFA: short-chain fatty acid
SIM: single ion monitoring
sOTU: sub-operational taxonomic unit
UniFrac: unique fraction metric

