## Supplemental Tables for "Characterization of gut microbiome and metabolome in *Helicobacter pylori* patients in an underprivileged community in the United States": H. pylori gut microbiome paper White et al Table S1.docx

**Table S1. Volcano plot data with all HELICO data set.**

| **Biochemical Name** | **FC** | **Log2(FC)** | ***p*-value** | **-Log10(*p*)** |
| --- | --- | --- | --- | --- |
| docosahexaenoic acid (22:6n3) | 1069.29 | 10.06 | 1.74E-01 | 0.76 |
| eicosapentaenoic acid (20:5n3) | 13.02 | 3.70 | 2.97E-01 | 0.53 |
| cis-13-16-docosadienoic acid (22:2n6) | 10.35 | 3.37 | 4.84E-02 | 1.31 |
| adrenic acid (22:4n6) | 9.17 | 3.20 | 3.81E-03 | 2.42 |
| docosapentaenoic acid (22:5n3) | 6.01 | 2.59 | 5.71E-03 | 2.24 |
| eicosatetraenoic acid (20:4n3) | 5.63 | 2.49 | 2.56E-01 | 0.59 |
| dihomo-gamma-linolenic acid (20:3n6) | 4.41 | 2.14 | 3.31E-02 | 1.48 |
| erucic acid (22:1n9) | 2.72 | 1.44 | 1.68E-01 | 0.77 |
| vaccenic acid (18:1n7) | 2.09 | 1.06 | 2.40E-01 | 0.62 |
| alpha-linolenic acid (18:3n3) | 2.06 | 1.04 | 2.88E-01 | 0.54 |
| nervonic acid (24:1n9) | 1.96 | 0.97 | 1.46E-01 | 0.83 |
| palmitic acid (16:0) | 1.94 | 0.96 | 7.98E-02 | 1.10 |
| palmitoleic acid (16:1n7) | 1.87 | 0.91 | 3.94E-01 | 0.40 |
| margaric acid (17:0) | 1.87 | 0.90 | 1.65E-01 | 0.78 |
| linoleic acid (18:2n6) | 1.65 | 0.73 | 5.31E-01 | 0.27 |
| myristic acid (14:0) | 1.59 | 0.67 | 3.99E-01 | 0.40 |
| arachidonic acid (20:4n6) | 1.46 | 0.54 | 1.03E-01 | 0.99 |
| cis-11,14-eicosadienoic acid (20:2n6) | 1.44 | 0.52 | 4.90E-01 | 0.31 |
| arachidic acid (20:0) | 1.32 | 0.40 | 4.95E-01 | 0.31 |
| stearidonic acid (18:4n3) | 1.29 | 0.36 | 8.52E-01 | 6.96E-02 |
| cis-11-eicosaenoic acid (20:1n9) | 1.26 | 0.34 | 6.59E-01 | 0.18 |
| myristoleic acid (14:1n5) | 1.25 | 0.32 | 7.72E-01 | 0.11 |
| behenic acid (22:0) | 1.06 | 9.03E-02 | 9.55E-01 | 2.01E-02 |
| stearic acid (18:0) | 1.05 | 6.62E-02 | 9.79E-01 | 9.28E-03 |
| pentadecanoic acid (15:0) | 1.04 | 5.54E-02 | 9.92E-01 | 3.51E-03 |
| oleic acid (18:1n9) | 0.98 | -2.39E-02 | 9.01E-01 | 4.52E-02 |
| lignoceric acid (24:0) | 0.93 | -0.11 | 7.22E-01 | 0.14 |
| osbond acid (22:5n6) | 0.54 | -0.88 | 3.57E-01 | 0.45 |
| gamma-linolenic acid (18:3n6) | 3.61E-02 | -4.79 | 7.57E-02 | 1.12 |
| mead acid (20:3n9) | 3.20E-02 | -4.97 | 3.39E-01 | 0.47 |
