## Supplemental Tables for "Characterization of gut microbiome and metabolome in *Helicobacter pylori* patients in an underprivileged community in the United States": H. pylori gut microbiome paper White et al Table S2.docx

**Table S2. Volcano plot data with High Bacteroidetes.**

| **Biochemical Name** | **FC** | **Log2(FC)** | ***p*-value** | **-Log10(*p*)** |
| --- | --- | --- | --- | --- |
| docosahexaenoic acid (22:6n3) | 2318.10 | 11.18 | 2.67E-01 | 0.57 |
| eicosapentaenoic acid (20:5n3) | 30.72 | 4.94 | 3.37E-01 | 0.47 |
| eicosatetraenoic acid (20:4n3) | 11.40 | 3.51 | 3.28E-01 | 0.48 |
| adrenic acid (22:4n6) | 9.33 | 3.22 | 1.01E-01 | 0.99 |
| docosapentaenoic acid (22:5n3) | 6.63 | 2.73 | 1.33E-01 | 0.88 |
| dihomo-gamma-linolenic acid (20:3n6) | 5.89 | 2.56 | 2.15E-01 | 0.67 |
| cis-13-16-docosadienoic acid (22:2n6) | 3.94 | 1.98 | 3.31E-01 | 0.48 |
| stearidonic acid (18:4n3) | 2.57 | 1.36 | 5.71E-01 | 0.24 |
| arachidonic acid (20:4n6) | 1.73 | 0.79 | 1.36E-01 | 0.87 |
| alpha-linolenic acid (18:3n3) | 1.65 | 0.72 | 5.73E-01 | 0.24 |
| erucic acid (22:1n9) | 1.31 | 0.39 | 6.61E-01 | 0.18 |
| palmitic acid (16:0) | 1.19 | 0.25 | 7.18E-01 | 0.14 |
| myristoleic acid (14:1n5) | 1.07 | 0.10 | 9.17E-01 | 3.78E-02 |
| cis-11,14-eicosadienoic acid (20:2n6) | 1.00 | -7.00E-03 | 9.93E-01 | 3.18E-03 |
| vaccenic acid (18:1n7) | 0.92 | -0.12 | 8.49E-01 | 7.09E-02 |
| margaric acid (17:0) | 0.92 | -0.12 | 8.39E-01 | 7.61E-02 |
| nervonic acid (24:1n9) | 0.84 | -0.26 | 5.94E-01 | 0.23 |
| linoleic acid (18:2n6) | 0.79 | -0.33 | 6.39E-01 | 0.19 |
| myristic acid (14:0) | 0.74 | -0.44 | 5.70E-01 | 0.24 |
| palmitoleic acid (16:1n7) | 0.72 | -0.48 | 3.58E-01 | 0.45 |
| arachidic acid (20:0) | 0.71 | -0.50 | 4.53E-01 | 0.34 |
| stearic acid (18:0) | 0.65 | -0.61 | 4.22E-01 | 0.37 |
| cis-11-eicosaenoic acid (20:1n9) | 0.42 | -1.24 | 2.24E-02 | 1.65 |
| pentadecanoic acid (15:0) | 0.40 | -1.34 | 4.52E-02 | 1.35 |
| behenic acid (22:0) | 0.35 | -1.50 | 7.07E-03 | 2.15 |
| lignoceric acid (24:0) | 0.30 | -1.73 | 4.83E-03 | 2.32 |
| oleic acid (18:1n9) | 0.30 | -1.74 | 3.35E-02 | 1.48 |
| osbond acid (22:5n6) | 0.20 | -2.33 | 1.08E-01 | 0.97 |
| gamma-linolenic acid (18:3n6) | 8.78E-02 | -3.51 | 9.36E-02 | 1.03 |
| mead acid (20:3n9) | 3.20E-02 | -4.97 | 3.39E-01 | 0.47 |
