## Supplemental Tables for "Characterization of gut microbiome and metabolome in *Helicobacter pylori* patients in an underprivileged community in the United States": H. pylori gut microbiome paper White et al Table S3.docx

**Table S3. Volcano plot data with High Firmicutes.**

| **Biochemical Name** | **FC** | **Log2(FC)** | ***p*-value** | **-Log10(*p*)** |
| --- | --- | --- | --- | --- |
| cis-13-16-docosadienoic acid (22:2n6) | 35.81 | 5.16 | 6.32E-02 | 1.20 |
| erucic acid (22:1n9) | 7.42 | 2.89 | 2.67E-01 | 0.57 |
| nervonic acid (24:1n9) | 4.27 | 2.09 | 2.35E-01 | 0.63 |
| alpha-linolenic acid (18:3n3) | 4.20 | 2.07 | 4.21E-01 | 0.38 |
| docosapentaenoic acid (22:5n3) | 4.20 | 2.07 | 1.89E-01 | 0.72 |
| myristic acid (14:0) | 2.89 | 1.53 | 4.32E-01 | 0.36 |
| margaric acid (17:0) | 2.71 | 1.44 | 3.74E-01 | 0.43 |
| cis-11-eicosaenoic acid (20:1n9) | 2.55 | 1.35 | 2.99E-01 | 0.52 |
| palmitic acid (16:0) | 2.52 | 1.33 | 2.80E-01 | 0.55 |
| behenic acid (22:0) | 2.48 | 1.31 | 1.69E-01 | 0.77 |
| vaccenic acid (18:1n7) | 2.44 | 1.29 | 2.41E-01 | 0.62 |
| adrenic acid (22:4n6) | 2.37 | 1.24 | 2.26E-01 | 0.65 |
| pentadecanoic acid (15:0) | 2.36 | 1.24 | 3.58E-01 | 0.45 |
| lignoceric acid (24:0) | 2.34 | 1.22 | 1.61E-01 | 0.79 |
| arachidic acid (20:0) | 2.29 | 1.19 | 2.08E-01 | 0.68 |
| myristoleic acid (14:1n5) | 2.12 | 1.08 | 6.25E-01 | 0.20 |
| cis-11,14-eicosadienoic acid (20:2n6) | 1.95 | 0.97 | 6.09E-01 | 0.22 |
| stearic acid (18:0) | 1.75 | 0.81 | 4.73E-01 | 0.33 |
| oleic acid (18:1n9) | 1.72 | 0.78 | 3.68E-01 | 0.43 |
| linoleic acid (18:2n6) | 1.40 | 0.48 | 6.50E-01 | 0.19 |
| dihomo-gamma-linolenic acid (20:3n6) | 1.37 | 0.45 | 4.87E-01 | 0.31 |
| palmitoleic acid (16:1n7) | 1.31 | 0.39 | 7.16E-01 | 0.15 |
| arachidonic acid (20:4n6) | 1.21 | 0.28 | 7.72E-01 | 0.11 |
| docosahexaenoic acid (22:6n3) | 1.00 | -3.20E-16 | 6.87E-03 | 2.16 |
| osbond acid (22:5n6) | 0.72 | -0.47 | 6.21E-01 | 0.21 |
| mead acid (20:3n9) | 3.20E-02 | -4.97 | 3.39E-01 | 0.47 |
| stearidonic acid (18:4n3) | 2.23E-02 | -5.49 | 1.76E-01 | 0.76 |
| eicosapentaenoic acid (20:5n3) | 1.12E-02 | -6.48 | 1.68E-01 | 0.77 |
| eicosatetraenoic acid (20:4n3) | 9.13E-03 | -6.78 | 1.18E-02 | 1.93 |
| gamma-linolenic acid (18:3n6) | 9.37E-05 | -13.38 | 6.70E-02 | 1.17 |
